# Gender disparity in computational biology research publications

**DOI:** 10.1101/070631

**Authors:** Kevin S. Bonham, Melanie I. Stefan

## Abstract

While women are generally underrepresented in STEM fields, there are noticeable differences between fields. For instance, the gender ratio in biology is more balanced than in computer science. We were interested in how this difference is reflected in the interdisciplinary field of computational/quantitative biology. To this end, we examined the proportion of female authors in publications from the PubMed and arXiv databases. There are fewer female authors on research papers in computational biology, as compared to biology in general. This is true across authorship position, year, and journal impact factor. A comparison with arXiv shows that quantitative biology papers have a higher ratio of female authors than computer science papers, placing computational biology in between its two parent fields in terms of gender representation. Both in biology and in computational biology, a female last author increases the probability of other authors on the paper being female, pointing to a potential role of female PIs in influencing the gender balance.

## Introduction

There is ample literature on the underrepresentation of women in STEM fields and the biases contributing to it. Those biases, though often subtle, are pervasive in several ways: they are often held and perpetuated by both men and women, and they are apparent across all aspects of academic and scientific practice. Undergraduate students show bias in favor of men both when rating their peers [4] and their professors [5]. Professors, in turn, are more likely to respond to e-mail from prospective students who are male [6]. They also show gender bias when hiring staff and deciding on a starting salary [7].

When looking at research output in the form of publication and impact, the story is complex: Women tend to publish less than men [8], are underrepresented in the more prestigious first and last author positions, and publish fewer single-author papers [9]. In mathematics, women tend to publish in lower-impact journals [1], while in engineering, women publish in journals with higher impact factors [2]. In general, however, articles authored by women are cited less frequently than articles authoreed by men [?,8], which might in part be due to men citing their own work more often than women do [10]. Inferring bias in these studies is difficult, since the cause of the disparity between male and female authorship cannot be readily determined. At the same time, when stories of scientific discoveries are told, gender biases are readily identified: Work by female scientists is more likely to be attributed to a male colleague [11], and biographies of successful female scientists perpetuate gender stereotypes [12]. Finally, the way in which evidence for gender bias is received is in itself biased: Male scientists are less likely to accept studies that point to the existence of gender bias than are their female colleagues [13].

Although gender imbalance seems to be universal across all aspects of the scientific enterprise, there are also more nuanced effects. In particular, not all disciplines are equally affected. For instance, in the biosciences over half of PhD recipients are now women, while in computer science, it is less than 20% [14]. This raises an intriguing question, namely how do the effects of gender persist in interdisciplinary fields where the parent fields are discordant for female representation?

To this end, we are interested in the gender balance in computational biology and how it compares to other areas of biology, since computational biology is a relatively young field at the disciplinary intersection between biology and computer science. We examined authorship on papers from Pubmed published between 1997 and 2014 and compared computational biology to biology in general. We found that in computational biology, there is a smaller proportion of female authors overall, and a lower proportion of female authors in first and last authorship positions than in all biological fields combined. This is true across all years, though the gender gap has been narrowing, both in computational biology and in biology overall. A comparison to computer science papers shows that computational biology stands between biology and computer science in terms of gender equality.

## Results and Discussion

In order to determine if there is a difference in the gender of authors in computational biology compared to biology as a whole, we used data from Pubmed, a database of biology and biomedical publications administered by the US National Library of Medicine. Pubmed uses Medical Subject Heading (MeSH) terms to classify individual papers by subject. The MeSH term “Computational Biology” is a subset of “Biology” and was introduced in 1997, so we restricted our analysis to primary articles published after this date (see S1 Fig A-B, Materials and Methods).

To determine the gender of authors, we used the web service Gender-API.com, which curates a database of first names and associated genders from government records as well as social media profiles. Gender-API searches provide information on the likely gender as well as confidence in the estimate based on the number of times a name appears in the database. We used bootstrap analysis to estimate the probability (*P_female_*) that an author in a particular dataset is female as well as a 95% confidence interval (see Materials and Methods).

We validated this method by comparing it to a set of 2155 known author:gender pairs from the biomedical literature provided by Filardo et. al. [15] Filardo and colleagues manually determined the genders of the first authors for over 3000 papers by searching for authors’ photographs on institutional web pages or social media profiles like LinkedIn. We compared the results obtained from our method of computational inference of gender for a subset of this data (see Materials and Methods), to the known gender composition of this author set. Infering author gender using Gender-API data suggested that *P_female_* = 0.373 ± 0.023 (Supplementary Fig 1C, black bar). Because the actual gender of each of these authors is known, we could also calculate the actual *P_female_* Using the same bootstrap method on actual gender (known female authors were assigned *P_female_* = 1, known male authors were assigned *P_female_* = 0), we determined that the real *P_female_* = 0.360 ± 0.018 (S1 Fig C, white bar).

Unfortunately, 43% of names used to query to Gender-API did not have associated gender information. These names, representing 26.6% of authors, were therefore excluded from our analysis. In order to ensure that this was not systematically skewing our results, we also determined the *P*_*female*_ in Filardo et al.’s known gender dataset excluding those authors with names that were not associated with a Gender-API record, giving *P_female_* = 0.381 ±.027 (S1 Fig C, white bar). Together, these results suggest that our method of automatically assigning gender using Gender-API gives comparable results to human-validated gender assignment, and that excluding names without clear gender information does not lead us to underestimate the proportion of women in our dataset.

We began our investigation of the gender make-up in biology and computational biology publications by analyzing the gender representation in primary publications from 1997 to 2014. Consistent with previous publications, women were substantially less likely to be in senior author positions than first author positions in publications labeled with the Biology (Bio) MeSH term (Last author, *P_female_* = 0.245 ± 0.002, First author, *P_female_* = 0.376 ± 0.003 (Fig 1A, Table 1). We observed the same trend in papers labeled with the computational biology (comp) MeSH term, though the *P_female_* at every author position was 4-6 percentage points lower. An analysis of publications by year suggests that the gender gaps in both biology and computational biology are narrowing, but by less than 1 percentage point per year (for bio, change in *P*_*female*_ = 0.0035 ± 0.0005/year, for comp, change in *P_female_* = 0.0049 ± 0.0008/*year*). However, the discrepancy between biology and computational biology has been consistent over time (Fig 1B).

**Fig 1.**
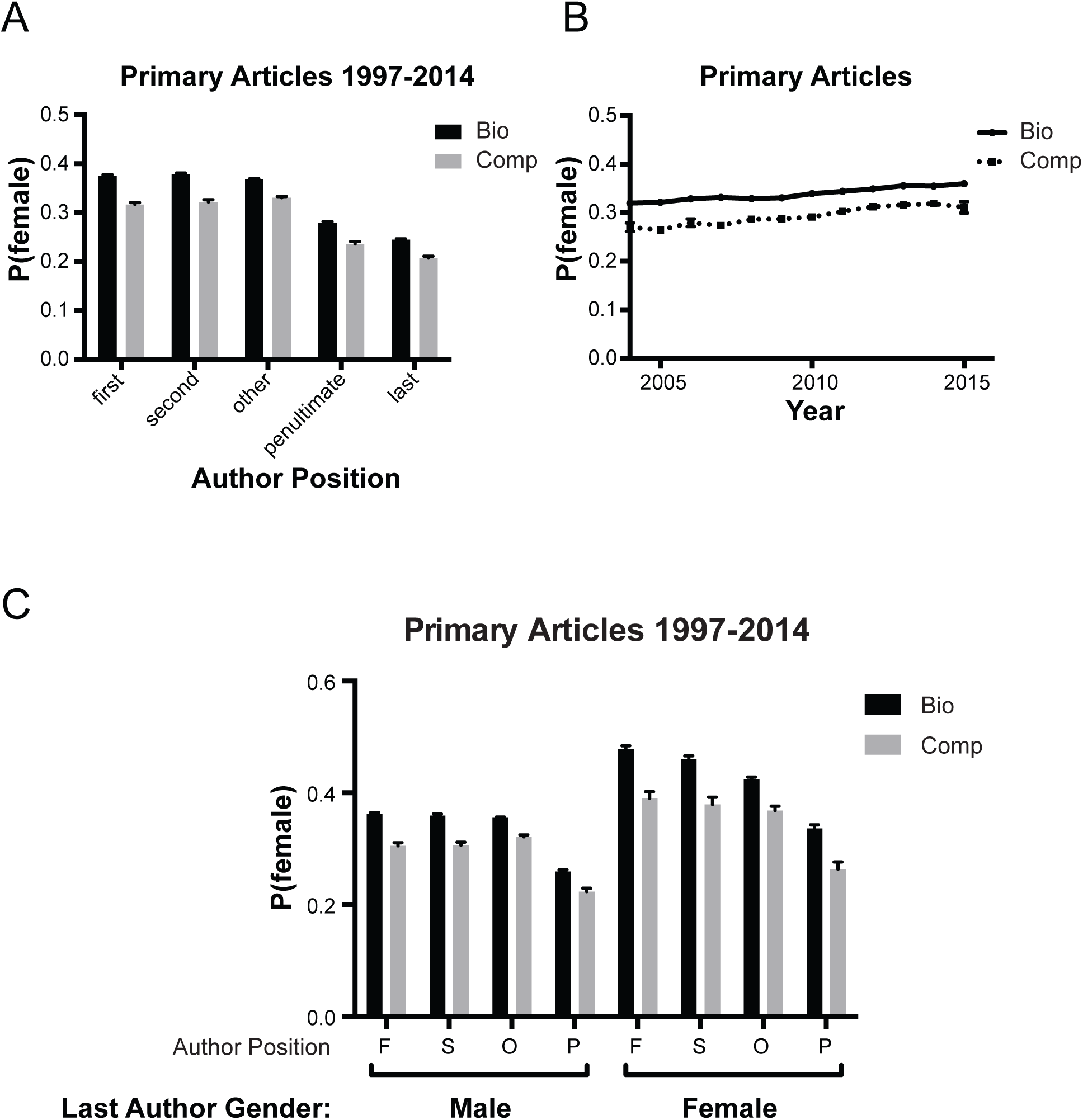
A: Mean probability that an author in a given position is female for primary articles indexed in Pubmed with the MeSH term Biology (black) or Computational Biology (grey). The bio dataset is inclusive of papers in the comp dataset. Error bars represent 95% confidence intervals. B: Mean probability that an author in a given position is female for primary articles indexed in Pubmed with the MeSH term Biology (black) or Computational Biology (grey). The bio dataset is inclusive of papers in the comp dataset. Error bars represent 95% confidence intervals. C: Mean probability that an author is female for publications in a given year. Error bars represent 95% confidence intervals. D: Mean probability that the first (F), second (S), penultimate (P) or other (O) author is female for publications where the last author is male (*P_female_* < 0.2) or female (*P_female_* > 0.8). Papers where the gender of the last author was uncertain or could not be determined were excluded. Error bars represent 95% confidence intervals.

**Table 1.** Proportion of Female Authors

| Dataset | Position | Mean | 95% CI |  |
| --- | --- | --- | --- | --- |
|  |  |  | lower | upper |
| bio | first | 0.376 | 0.373 | 0.378 |
|  | second | 0.379 | 0.376 | 0.381 |
|  | other | 0.368 | 0.367 | 0.370 |
|  | penultimate | 0.279 | 0.277 | 0.282 |
|  | last | 0.245 | 0.243 | 0.247 |
| comp | first | 0.316 | 0.312 | 0.320 |
|  | second | 0.322 | 0.317 | 0.327 |
|  | other | 0.331 | 0.328 | 0.333 |
|  | penultimate | 0.236 | 0.231 | 0.241 |
|  | last | 0.207 | 0.203 | 0.211 |

One possible explanation for the difference in male and female authorship position might be a difference in role models or mentors. If true, we would expect studies with a female principal investigator to be more likely to attract female collaborators. Conventionally in biology, the last author on a publication is the principal investigator on the project. Therefore, we looked at two subsets of our data: publications with a female last author (*P_female_* > 0.8) and those with a male last author (*P_female_* < 0.2). We found that women were substantially more likely to be authors at every other position if the paper had a female last author than if the last author was male (Fig 1C, Table 2). It is possible that female trainees are be more likely to pursue computational biology if they have a mentor that is also female. Since women are less likely to be senior authors, this might reduce the proportion of women overall. However, we cannot determine if the effect we observe is instead due to a tendancy for women that pursue computational biology to select female mentors.

**Table 2.** Proportion of Female Authors with Female PI

| Dataset | Position | Male Last Author |  |  | Female Last Author |  |  |
| --- | --- | --- | --- | --- | --- | --- | --- |
|  |  | Mean | 95% CI |  | Mean | 95% CI |  |
|  |  |  | lower | upper |  | lower | upper |
| bio | first | 0.362 | 0.359 | 0.365 | 0.478 | 0.472 | 0.484 |
|  | second | 0.359 | 0.357 | 0.362 | 0.460 | 0.454 | 0.466 |
|  | other | 0.355 | 0.353 | 0.357 | 0.425 | 0.421 | 0.428 |
|  | penultimate | 0.259 | 0.256 | 0.263 | 0.336 | 0.330 | 0.343 |
| comp | first | 0.305 | 0.300 | 0.311 | 0.390 | 0.378 | 0.402 |
|  | second | 0.306 | 0.300 | 0.312 | 0.379 | 0.366 | 0.392 |
|  | other | 0.321 | 0.318 | 0.324 | 0.368 | 0.361 | 0.376 |
|  | penultimate | 0.223 | 0.218 | 0.229 | 0.263 | 0.249 | 0.277 |

Though MeSH terms enable sorting a large number of papers regardless of where they are published, the assignment of these terms is a manual process and may not be comprehensive for all publications. As another way to qualitatively examine gender differences in publishing, we examined different journals, since some journals specialize in computational papers, while others are more general. We looked at the 123 journals that had at least 1000 authors in our bio dataset, and determined *P_female_* for each journal separately (Fig 2A). Of these journals, 21 (14%) have titles indicative of computational biology or bioinformatics, and these journals have substantially lower representation of female authors. The 3 journals with the lowest female representation and 6 out of the bottom 10 are all journals focused on studies using computational methods. Only 4 computational biology/bioinformatics journals are above the median of female representation.

**Fig 2.**
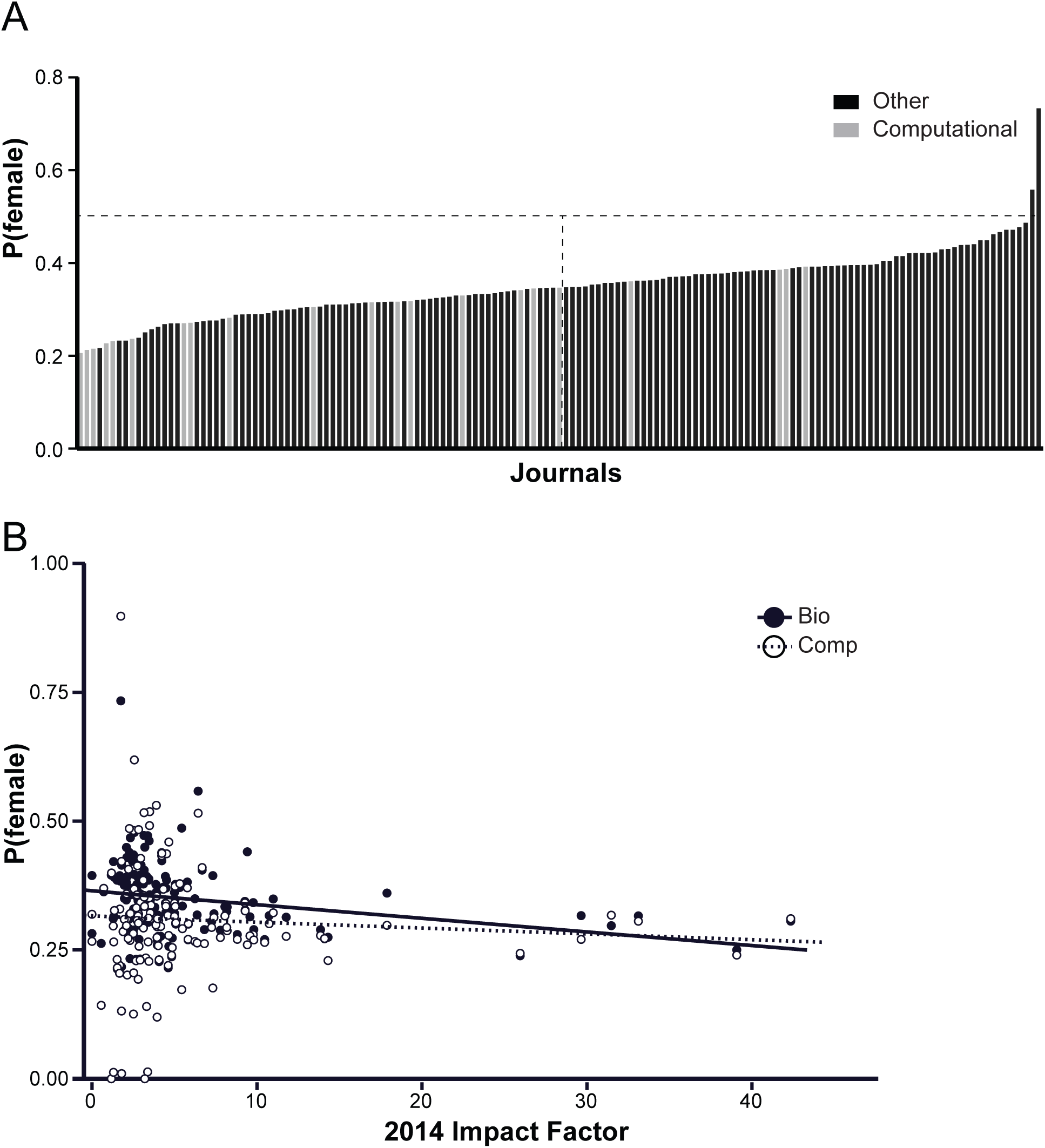
A: Mean probability that an author is female for every journal that had at least 1000 authors in our dataset. Grey bars represent journals that have the words “Bioinformatics,” “Computational,” “Computer,” “System(s),” or “-omic(s)” in their title. Vertical line represents the median for female author representation. See also S1 Table. B: Mean probability that an author is female for articles in the “Bio” dataset (black dot) or in the “Comp” dataset (open square) for each journal that had at least 1000 authors plotted against the journals’ 2014 impact factor. Journals that had computational biology articles are included in both datasets. An ordinary least squares regression was performed for each dataset. Bio: *m* = -0.00264, *PZ*>|z| = 0.0022. Comp: *m* = -0.000789, *PZ*>|z| = 0.568.

One possible explanation might be that women are less likely to publish in high-impact journals, so we considered the possibility that the differences in the gender of authors that we observe could be the result of differences in impact factor between papers published in biology versus computational biology publications. We compared the *P_female_* of authors in each journal with that journal’s 2014 impact factor (Fig 2B). There is a marginal but significant negative correlation (-0.00264, *PZ*>|z| = 0.0022) between impact factor and gender for the biology dataset. This is in contrast to previous studies from engineering that have found that women tend to publish in higher-impact journals [2]. It is, however, consistent with a previous studies from mathematics [1]. By contrast, there is no significant correlation (*PZ*>|z| = 0.568) between impact factor and *P_female_* in computational biology publications. Further, for journals that have articles labeled with the computational biology MeSH term, the *P_female_* for those articles is the same or lower than that for all biology publications in the same journal.

We also examined whether computational biology or biology articles tend to have higher impact factors. Bootstrap analysis of authors in each dataset suggest that computational biology publications tend to be published in journals with a higher impact factor (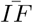 = 7.25 ± 0.04) than publications in biology as a whole (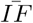 = 6.5 ± 0.02). However, given the magnitude of the correlation between IF and *P_female_*, this difference is unlikely to explain the differences in *P_female_* observed between our computational biology and biology datasets. Taken together, these data suggest that the authors of computational biology papers are less likely to be women than the authors of biology papers generally.

We turned next to an investigation of biological fields relative to computer science. Since Pubmed does not index computer science publications, we cannot compare the computational biology dataset to computer science research papers directly. Instead, we investigated the gender balance of authors of manuscripts submitted to arXiv, a preprint repository for academic papers used frequently by quantitative fields like mathematics and physics. These preprint records cannot be compared to peer-reviewed publications indexed on pubmed, but a “quantitative biology” (qb) section was added to arXiv in 2003. Quantitative biology is not necessarily equivalent to computational biology, and analysis of arXiv-qb papers that have been published and indexed on pubmed suggests that only a fraction of them are labeled with the “computational biology” MeSH term. However, this does allow us to make an apples-to-apples comparision between a field of biology and computer science. There are relatively few papers preprints prior to 2007, so we compared preprints in “quantitative biology” to those in “computer science” from 2007-2016.

Women were more likely to be authors in quantitative biology manuscripts than in computer science manuscripts in first, second, and middle author positions (Fig 3A, Table 3). We found no significant difference in the frequency of female authors in the last or penultimate author positions in these two datasets, though the conventions for determining author order are not necessarily the same in computer science as in biology. Nevertheless, women had higher representation in quantitative biology than in computer science for all years except 2009 (Fig 3B). Interestingly, there is a slight but significant (0.0052/*year, P*Z>|z| < 0.005) increase in the proportion of female authors over time in quantitative biology, while there’s no significant increase in female representation in computer science preprints.

**Fig 3.**
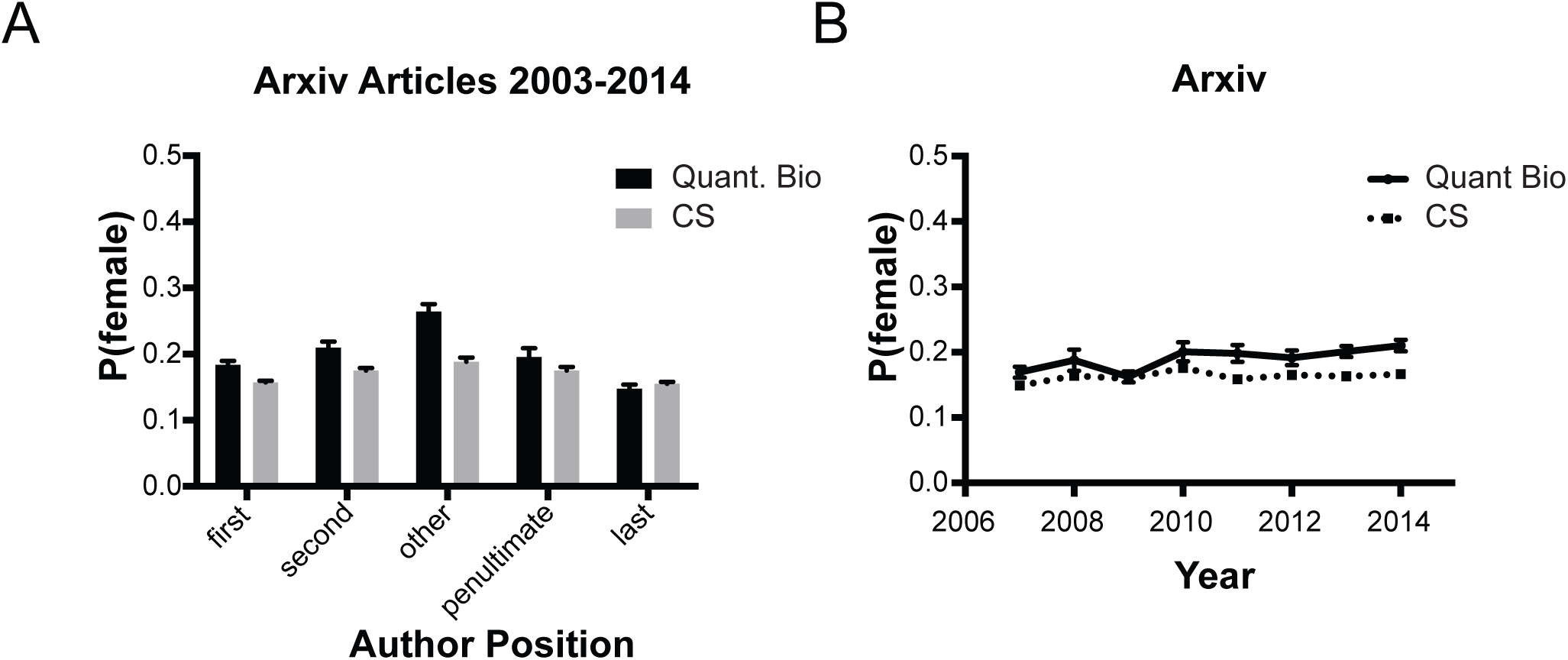
A: Mean probability that an author in a given position is female for all preprints in the arXiv quantitative biology (black) or computer science (grey) categories between 2007 and 2014. Error bars represent 95% confidence intervals. B: Mean probability of authors being female in arXiv preprints in a given year. Error bars represent 95% confidence intervals. Slopes were determined using ordinary least squares regression. The slope for q-bio is slightly positive (p < 0.05), but the slope for cs is not.

**Table 3.** Proportion of Female Authors in Arxiv

| Dataset | Position | Mean | 95% CI |  |
| --- | --- | --- | --- | --- |
|  |  |  | lower | upper |
| arxivbio | first | 0.184 | 0.178 | 0.190 |
|  | second | 0.210 | 0.200 | 0.219 |
|  | other | 0.265 | 0.253 | 0.276 |
|  | penultimate | 0.196 | 0.183 | 0.209 |
|  | last | 0.148 | 0.141 | 0.155 |
| arxivcs | first | 0.157 | 0.155 | 0.160 |
|  | second | 0.175 | 0.172 | 0.179 |
|  | other | 0.188 | 0.182 | 0.195 |
|  | penultimate | 0.175 | 0.170 | 0.181 |
|  | last | 0.155 | 0.153 | 0.158 |

Taken together, our results suggest that computational biology lies between biology in general and computer science when it comes to gender representation in publications. This is perhaps not surprising given the interdisciplinary nature of computational biology. Compared to biology in general, computational biology papers have fewer female authors, and this is consistent across all authorship positions. Importantly, this difference is not due to a difference in impact factor between computational biology and general biology papers.

Articles with a female last author tend to have more female authors in other positions and this is true for both biology in general and computational biology. Since the last author position is most often occupied by the principal investigator of the study, this suggests that having a woman as principal investigator has a positive influence on the participation of women. This resonates with findings by Macaluso et al., who studied the nature of authorship contribution by gender in PLoS publications [3]. They found that if the corresponding author of a paper was female, then there was also a greater proportion of women across almost all authorship roles (data analysis, experimental design, performing experiments, and writing the paper). In contrast, if the corresponding author was male, then men were dominating all authorship roles except for performing experiments, which remained female-dominated. The reasons for this are difficult to ascertain. It could be the case that female PIs tend to work in more female-dominated sub-fields and therefore naturally have more female co-authors. It is also possible that female PIs are more likely to recognise contributions by female staff members, or that they are more likely to attract female co-workers and collaborators. Our publication data cannot differentiate between those two (and other) explanations, but points to the important role that women in senior positions may play as role models for trainees.

Since biology attracts more women than computer science, we suspect that many women initially decide to study biology and later become interested in computational biology. If this is the case, understanding what factors influence the field of study will provide useful insight when designing interventions to help narrow the gender gap in computer science and computational biology.

## Materials and Methods

### Datasets

#### Biology publications 1997-2014 (bio)

This dataset [16] contains all English language publications under the MeSH term “Biology” published between 1997 and 2014, excluding many non-primary sources. This set contains 204,767 records. Downloaded 12 February, 2016. Search term: ("Biology"[Mesh]) NOT (Review[ptyp] OR Comment[ptyp] OR Editorial[ptyp] OR Letter[ptyp] OR Case Reports[ptyp] OR News[ptyp] OR “Biography” [Publication Type]) AND ("1997/01/01"[PDAT]: “2014/12/31”[PDAT]) AND english[language]Same as above [16], except using MeSH term “Computational Biology”. Only uses papers where this is a major term. Date range was selected because this MeSH term was introduced in 1997. This dataset is a subset of the “bio” dataset (all of the papers in this dataset are contained within “bio") and contains 43,198 records. Downloaded 12 February, 2016. Search term: ("Computational Biology"[Majr]) NOT (Review[ptyp] OR Comment[ptyp] OR Editorial[ptyp] OR Letter[ptyp] OR Case Reports[ptyp] OR News[ptyp] OR “Biography” [Publication Type]) AND ("1997/01/01"[PDAT]: “2014/12/31”[PDAT]) AND english[language]

#### Medical Papers

Subset of author and gender data from Filardo et.al [15]. This dataset did not contain author first names or unique publication identifiers. We searched pubmed for the title, author and publication date, and were able to identify 2155/3153 publications to analyze. Publications with no matching search results or with multiple matching search results were excluded.

#### arXiv Quantitative Biology (q-bio)

This dataset [17] contains all preprints with the label “q-bio” from 2003 (when the section was introduced) to 2014. This set contains 41,637 records and was downloaded on 10 June, 2016.

#### arXiv CS (cs)

This dataset [17] contains all preprints with the label “cs” from 2003 to 2014, and contains 188,617 records. Downloaded on 10 June, 2016. There are 1412 preprints that are found in both the qbio and cs dataset (3.4% of bio and 0.75% of cs).

### Gender Inference

Genders were determined using Gender-API (http://gender-api.com), which compares first names to a database compiled from government sources as well as from crawling social media profiles and returns a gender probability and a measure of confidence based on the number of times the name appears in the database. The API was queried with the 74,760 unique first names in the dataset (24 May, 2016).

Mean gender probabilities were determined using bootstrap analysis. Briefly, for each dataset, authors were randomly sampled with replacement to generate a new dataset of the same size. The mean *P*_*female*_ for each sample was determined excluding names for which no gender information was available ( 26.6% of authors). The reported *P*_*female*_ represents the mean of means for 1000 samples. Error bars in figures represent 95% confidence intervals. Code and further explanation can be found on github [18].

Author positions were assigned based on the number of total authors. In papers with 5 or more authors, all authors besides first, second, last and penultimate were designated “other.” Papers with 3 authors were assigned only first, second and last, papers with two authors were assigned only first and last, and single-author papers were assigned only first.

## Regression Analysis

We used ordinary least squares regression analyses on IF and *P_female_* using the the GLM.jl package for the julia programming language. Correlations were considered significant if *P*Z>|*z*|< 0.05

## Acknowledgements

The authors would like to thank Markus Perl for the free use of Gender-API - contactgender-api.com; Casper Strømgren for the free use of genderize.io infogenderize.io; Giovanni Filardo for sharing data [15]; Johanna Gutlerner, Marshall Thomas, Diane Lam, and other members of the Curriculum Fellows Program (CFP) at Harvard Medical School (HMS) for helpful feedback and discussions; and The HMS CFP and Educational Laboratory for resources and mentorship

## Supporting Information

**S1 Fig. A.**
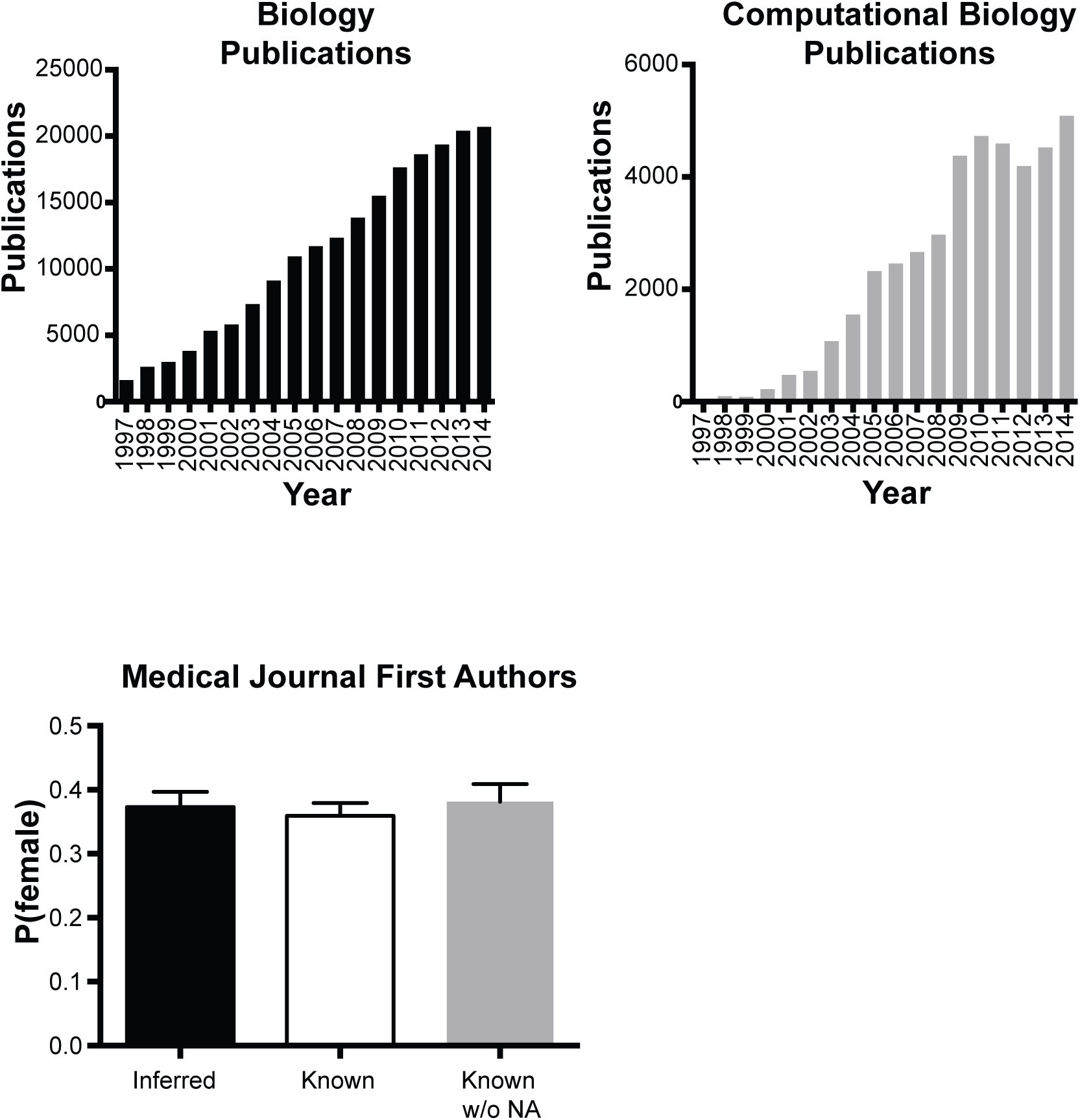
: Number of primary publications per year indexed under the “Biology” MeSH term. B: Number of primary publications per year indexed with “Computational Biology” as a major MeSH term. C: Comparison of computational gender inference (black) with known genders (white) for the dataset from Filardo et. al. [15]. Grey represents the known proportion of female authors when excluding names for which the gender could not be computationally inferred. Error bars represent 95% confidence intervals.

**S2 Fig. A:**
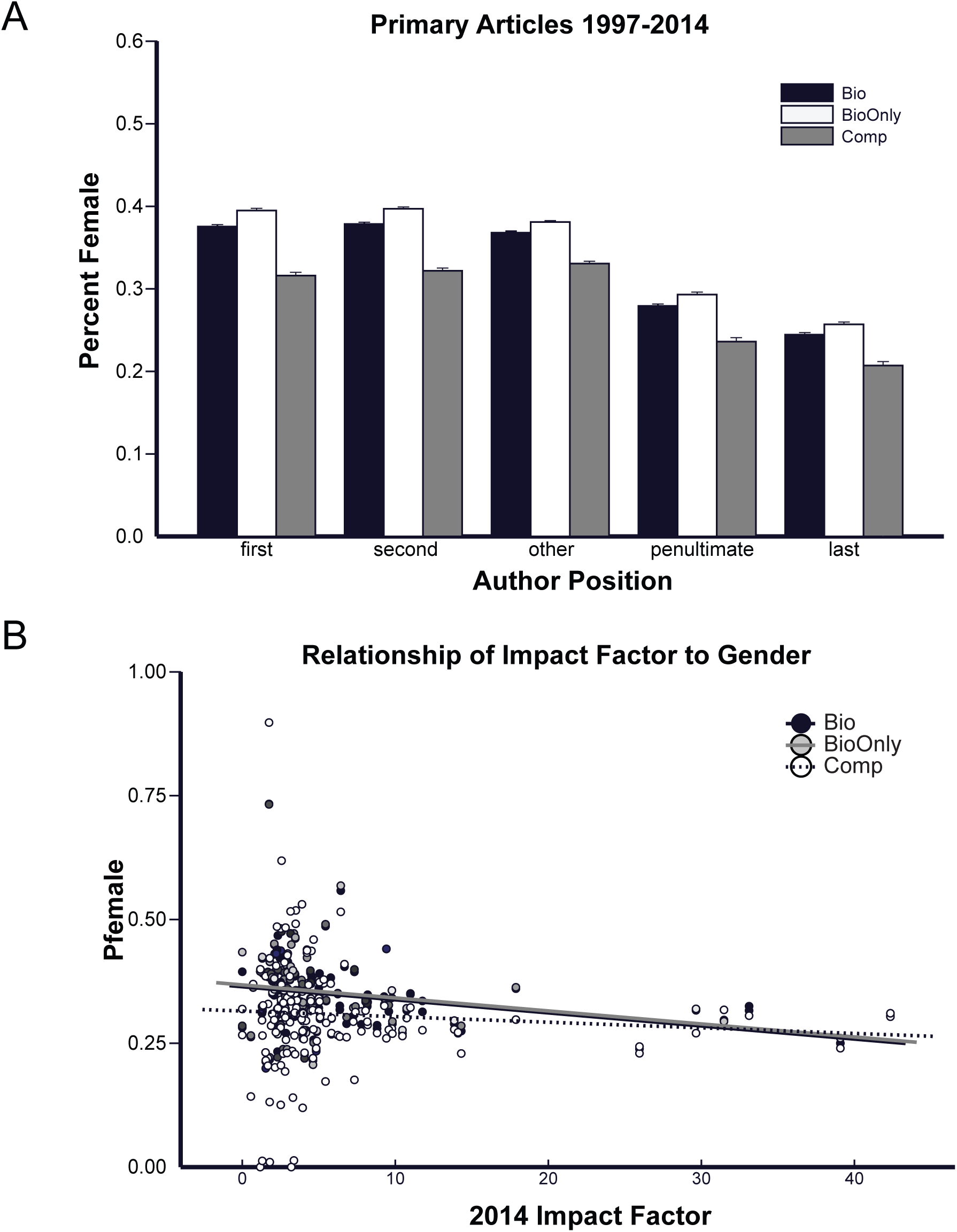
Mean probability that an author in a given position is female for primary articles indexed in Pubmed with the MeSH term Biology (black), Computational Biology (gray) or for those articles with Biology but not Computational biology (white). Error bars represent 95% confidence intervals. B: Mean probability that an author is female for articles in the “Bio” dataset (black) in the “Comp” dataset (white), or for articles in the Bio but not Comp (gray) for each journal that had at least 1000 authors plotted against the journals’ 2014 impact factor. Excluding computational publications from the biology dataset does not substantially alter the correlation between impact factor and *P_female_*

---

**S1 Table.***P*_*female*_ for each journal with at least 1000 authors in the bio dataset. Journals identified as primarily computational are shaded grey.

